# Depression Severity Modulates the Cognitive Process of Facial Affect Representation: Evidence from a Genetic-Algorithm Face Synthesis Task

**DOI:** 10.64898/2026.06.01.729421

**Authors:** Bliss H. Cui, Peter J. Bex

## Abstract

Depression is associated with biased facial emotion processing, yet existing paradigms measure only the recognition of pre-selected stimuli, conflating the process of forming an internal representation with its endpoint product. Here, we used a genetic-algorithm (GA) face synthesis task to disentangle these components. Fifty-seven undergraduates (40 Low Risk, 17 At Risk by Beck Depression Inventory-II) iteratively evolved photorealistic 3D faces to match their internal representations of 13 emotions across seven generations in a 199-dimensional face shape space. We extracted five evolutionary metrics capturing how participants constructed their representations (convergence speed, velocity, stability, range) and the structure they ultimately produced (peak intensity). At Risk participants converged on embarrassment representations significantly faster than Low Risk participants (*d* = 0.88, *p* = .001, FDR-corrected, permutation-validated), reaching their template in roughly half the number of generations. Anger representations showed greater evolutionary instability in the At Risk group (*d* = -0.75, *p* = .029, permutation-validated). Critically, endpoint face intensities did not differ between groups for any emotion. These results suggest that depression severity is associated with rigid self-conscious emotion schemas and unstable anger representations during face generation, reflecting altered cognitive processes rather than distorted perceptual products. The findings extend cognitive schema theory to the domain of facial affect representation and highlight self-conscious emotions as an underexplored locus of depression-related perceptual bias.

## 1. Introduction

### 1.1 Depression and Facial Emotion Processing

Depression fundamentally alters how people perceive emotional expressions in faces. A substantial body of evidence, spanning behavioral, neuroimaging, and electrophysiological methods, has documented systematic biases in facial emotion processing among individuals with depressive symptoms. These biases are thought to both maintain and exacerbate depressive episodes by distorting the social information that guides interpersonal behavior (Gotlib & Joormann, 2010).

Meta-analytic evidence confirms that individuals with major depressive disorder (MDD) show broad deficits in facial emotion recognition. Dalili et al. (2015) synthesized data from 22 independent samples and found a small but reliable overall impairment (*g* = -0.16), with the largest deficits emerging for happiness recognition. Demenescu et al. (2010) reached similar conclusions in an earlier meta-analysis, emphasizing that the pattern is not uniform across emotions. Notably, sadness recognition tends to be preserved or even enhanced in depression (Bourke et al., 2010), consistent with mood-congruent processing accounts. Happiness recognition, by contrast, is specifically impaired, suggesting that anhedonia may selectively erode sensitivity to positive social signals.

At the neural level, depression is associated with potentiated amygdala reactivity to negative emotional faces, even when stimuli are presented below the threshold of conscious awareness. Dannlowski et al. (2009) demonstrated automatic mood-congruent amygdala responses to masked sad and angry faces in currently depressed participants, implicating early, preattentive processing stages. Stuhrmann et al. (2011) reviewed the broader neuroimaging literature and identified a convergent pattern: hyperactivation of limbic regions to negative faces coupled with hypoactivation to positive faces, consistent with a negativity bias that operates across multiple levels of the processing hierarchy.

Despite this rich evidence base, a critical limitation pervades the existing literature. Virtually all studies of facial emotion processing in depression rely on recognition paradigms in which participants are shown pre-selected facial stimuli and asked to label, rate, or categorize them. These paradigms capture the endpoint product of perception (i.e., the accuracy or speed of a categorization decision) but reveal little about the underlying process through which an internal representation of an emotional face is constructed and refined.

### 1.2 Limitations of Recognition Paradigms

Recognition tasks present fixed stimuli and measure accuracy or reaction time, providing a snapshot of the final perceptual judgment. However, this approach has several limitations that constrain what can be inferred about the cognitive mechanisms underlying depression-related biases.

First, the stimuli in recognition paradigms are typically experimenter-selected, introducing potential selection bias. As Binetti et al. (2022) argued, the faces used in standard emotion recognition batteries represent one possible instance of each emotion category, not necessarily the prototype that any given participant would generate. Individual differences in what counts as a prototypical angry or sad face are substantial (Binetti et al., 2022) but are invisible to paradigms that hold the stimulus constant across participants.

Second, recognition tasks cannot distinguish between two fundamentally different cognitive mechanisms that could produce the same behavioral outcome. A depressed individual who is slower to recognize happiness could have an intact internal template for happy faces but an impaired decision process for accessing or labeling it, or alternatively could have a genuinely distorted internal template that deviates from the normative representation. These two possibilities have very different theoretical implications: the former implicates a post-perceptual labeling deficit, whereas the latter implicates a representational distortion at the level of the stored emotion schema. Recognition paradigms, by measuring only the output of the full processing chain, cannot adjudicate between these accounts.

Third, recognition paradigms are inherently constrained to the set of emotions for which standardized stimulus sets exist. The field has consequently focused heavily on a small number of canonical emotion categories, leaving self-conscious emotions such as embarrassment, shame, and pride largely unexplored in the context of depression and face perception. This gap is notable because self-conscious emotions are theoretically central to depressive phenomenology (Tangney & Dearing, 2002).

### 1.3 The Genetic-Algorithm Face Task

The genetic-algorithm (GA) face task offers a paradigm that overcomes these limitations by shifting control of the stimulus from the experimenter to the participant. Introduced by Carlisi et al. (2021) and expanded by Binetti et al. (2022), the GA face task asks participants to iteratively evolve photorealistic 3D faces to match their internal representation of each target emotion. Participants are presented with a population of randomly generated faces and select those that best match the target emotion. Selected faces contribute their features to the next generation through crossover, producing a new population that, on average, more closely approximates the participant’s internal template. This process repeats across multiple generations, producing an evolutionary trajectory that reveals how the representation is constructed over time.

In the present study, the faces are generated using the Basel Face Model (Paysan et al., 2009), a 199-dimensional shape space derived from principal component analysis of 3D face scans. Each face is a point in this high-dimensional space, and the genetic algorithm searches this space according to the participant’s selections and represents an evolutionary trajectory. The resulting trajectory provides a rich data source: it captures not only the endpoint face (the product) but also the dynamics of the search process, including how quickly the participant converges on a stable representation, how variable the search is across generations, and how rapidly the representation changes over time.

Prior work with the GA face task has revealed profound individual differences in facial emotion representation. Binetti et al. (2022) found that the internal templates participants evolved for the same emotion varied substantially across individuals, suggesting that emotion categories are represented with far more heterogeneity than standardized stimulus sets would imply. Carlisi et al. (2021) demonstrated the feasibility of the approach and validated it against expert-rated emotion categories. However, neither study examined how clinical or subclinical dimensions of psychopathology relate to the evolutionary dynamics of face construction.

The present study extends this paradigm in two critical ways. First, we expanded the emotion set to 13 categories, informed by Le Mau et al. (2021), who established these 13 emotion dimensions through standardized ratings of naturalistic emotional expressions from the Actors Acting corpus. This selection includes self-conscious emotions (embarrassment, shame, pride, contempt) and cognitive emotions (interest, awe, amusement) alongside basic emotions, providing broader coverage of the affective landscape than prior GA studies. Second, we introduced five evolutionary metrics that quantify distinct aspects of the face construction process: peak intensity, velocity of change, population stability, search range, and convergence speed. These metrics allow us to separately assess the product of face construction (what the participant ultimately produced) and the process by which they got there (how they searched the representational space).

### 1.4 Self-Conscious Emotions and Depression

Self-conscious emotions, including shame, guilt, embarrassment, and pride, occupy a distinctive position in the architecture of affective experience. Unlike basic emotions, which arise primarily in response to external events, self-conscious emotions require self-referential processing: they involve the evaluation of one’s own behavior, appearance, or character against internalized standards (Tracy & Robins, 2004). This self-evaluative component makes them theoretically central to depression, which is characterized by pervasive negative self-assessment and harsh self-judgment (Beck, 2019).

Shame proneness, in particular, shows a robust association with depressive symptomatology. Orth et al. (2006) found that trait shame correlated *r* = .57 with depression severity, with this relationship partially mediated by maladaptive rumination. The tendency to experience shame, to feel exposed and diminished before the real or imagined gaze of others, maps closely onto the core cognitive features of depression: global, stable, internal attributions for negative events (Abramson et al., 1989) and heightened sensitivity to social evaluation (Tangney & Dearing, 2002).

Embarrassment, one focus of the present investigation, is closely related to shame but is distinguished by its emphasis on public self-presentation rather than global self-worth. Keltner and Buswell (1997) proposed that embarrassment arises from the conjunction of two factors: a perceived deviation from personal or social standards and the awareness that this deviation is witnessed or evaluated by others. Both factors are amplified in depression. Depressed individuals exhibit elevated maladaptive perfectionism (Hewitt & Flett, 1991), perceive more frequent violations of their own standards, and show heightened sensitivity to the evaluative gaze of others (Orth et al., 2006). Embarrassment may therefore represent a particularly salient emotion category for individuals with elevated depressive symptoms.

Despite the theoretical centrality of self-conscious emotions to depression, they have been almost entirely absent from the facial emotion processing literature. The GA face task, by generating faces per individual in real time, rather than drawing from a fixed stimulus set, removes this constraint and allows self-conscious emotions to be studied within the same paradigm as basic emotions.

### 1.5 Anger and Depression

Although anger is not traditionally considered a hallmark of depression, accumulating evidence suggests it is a core rather than peripheral feature. Judd et al. (2013) found that 37% of patients with unipolar MDD reported clinically significant irritability and anger, and that anger attacks were associated with greater depression severity, more comorbidity, and worse functional outcomes.

The mechanism linking anger to depression may involve emotion dysregulation. Besharat et al. (2013) demonstrated that the relationship between anger and depressive symptoms is mediated by difficulties in emotion regulation, particularly the inability to effectively modulate the intensity and duration of anger episodes. Loch et al. (2024) extended this framework by showing that individuals with MDD exhibit more variable and less flexible emotion regulation strategy use compared to healthy controls, suggesting a broader pattern of regulatory instability rather than a fixed bias toward any single emotion.

This dysregulation framework has implications for how anger representations may be constructed in a generative task. If depression is associated with unstable anger regulation, this instability could manifest in the evolutionary search dynamics: more variable face populations across generations, reflecting oscillation between competing anger representations rather than smooth convergence on a stable template. This contrasts with the schema rigidity account for self-conscious emotions, raising the possibility that depression may have qualitatively different effects on different emotion categories.

### 1.6 Process Versus Product: A Schema-Theoretic Framework

Beck’s cognitive theory of depression (Beck, 2019) posits that depressive cognition is characterized by rigid, automatically activated negative schemas that bias information processing across multiple domains. These schemas are not merely beliefs but structured knowledge representations that guide perception, attention, memory, and interpretation. Disner et al. (2011) reviewed neuroimaging evidence supporting the schema model, showing that depression is associated with enhanced bottom-up processing of schema-congruent stimuli and impaired top-down cognitive control.

The GA face task provides a novel opportunity to examine schema theory in the domain of facial affect representation. Schema rigidity would account for faster convergence during the evolutionary search, because rigid schemas entail deploying a pre-formed template rather than openly exploring the face space. In other words, rigid schemas would compress the process of representation construction, leading to earlier convergence with less generational variability. This is a process-level account: it concerns how the representation is built, not what it ultimately looks like.

Conversely, emotion dysregulation would manifest as instability in the evolutionary trajectory: greater oscillation across generations, a failure to maintain a consistent representation from one generation to the next. This, too, is a process-level phenomenon. Neither schema rigidity nor dysregulation necessarily entails differences in the endpoint face (the product), because a rigid schema and a carefully explored representation could converge on the same final face, and an unstable search could nonetheless arrive at an average endpoint similar to that of a stable search.

The distinction between process and product is therefore central to interpreting any depression-related effects in this paradigm. If depression modulates the dynamics of face construction (how participants search the representational space) without altering the endpoint (what they ultimately produce), this would implicate the cognitive process of representation formation rather than the perceptual product itself.

### 1.7 The Present Study

The present study examined whether depression severity modulates the evolutionary dynamics of facial emotion construction in a GA face task. Undergraduates completed the Beck Depression Inventory-II (BDI-II; Beck et al., 1996) and were classified into Low Risk and At Risk groups based on the established clinical cutoff. They then completed the GA face task for 13 emotions spanning basic, self-conscious, and cognitive categories.

Five evolutionary metrics were extracted from each participant’s generational trajectory for each emotion, capturing both the process of face construction (convergence speed, velocity, stability, and range) and its product (peak intensity). Group comparisons were conducted with appropriate corrections for multiple comparisons and validated through permutation testing. Sensitivity analyses assessed the robustness of results to analytic threshold choices.

This study represents, to our knowledge, the first application of the GA face paradigm to a clinical dimension, and the first to examine self-conscious emotion representations in relation to depression severity using a generative face task.

## 2. Methods

### 2.1 Participants

Fifty-seven undergraduate students (43 female, 14 male; *M* age = 19.51 years, *SD* = 1.73, range = 18–30) were recruited from the Northeastern University psychology participant pool and compensated with course credit. All participants had normal or corrected-to-normal vision, by self-report, and provided written informed consent. The study protocol was approved by the Northeastern University Institutional Review Board. A subset of 53 participants also completed an Empathy Quotient (EQ) task reported separately.

Participants were classified into two groups based on their Beck Depression Inventory-II (BDI-II) total scores using the established clinical cutoff of 14 (Beck et al., 1996): Low Risk (BDI-II < 14; *n* = 40, 30 female; *M* age = 19.38, *SD* = 1.35) and At Risk (BDI-II ≥ 14; *n* = 17, 13 female; *M* age = 19.82, *SD* = 2.35). The groups did not differ in sex composition (χ² < 0.01, *p* = 1.000) or age (*t*(22.3) = -0.89, *p* = .376). Within the At Risk group, BDI-II severity ranged from mild (14–19; *n* = 11) to moderate (20–28; *n* = 4) to severe (29–63; *n* = 2), and four participants reported a prior clinical diagnosis of depression.

The term ’At Risk’ is used throughout to denote elevated depressive symptomatology as indexed by the BDI-II, a self-report screening instrument. This designation does not imply a clinical diagnosis of major depressive disorder. This subclinical approach is consistent with dimensional models of depression and has been widely used to study cognitive biases associated with depressive symptoms in nonclinical populations (Gotlib & Joormann, 2010).

### 2.2 Beck Depression Inventory-II

The BDI-II (Beck et al., 1996) is a 21-item self-report questionnaire assessing the severity of depressive symptoms over the past two weeks. Each item is rated on a 0–3 scale, yielding total scores from 0 to 63. The BDI-II has strong psychometric properties, with high internal consistency (α = .91 in outpatient samples; Beck et al., 1996) and well-established cutoff scores: 0–13 (minimal), 14–19 (mild), 20–28 (moderate), and 29–63 (severe). In the present sample, the full-sample BDI-II distribution was: *M* = 10.42, *SD* = 7.44, Mdn = 9, range = 0–34. Low Risk participants scored *M* = 6.30, *SD* = 3.59, and At Risk participants scored *M* = 20.12, *SD* = 5.69.

### 2.3 Apparatus and Stimuli

Stimuli were presented on a 27-inch iMac (5120 × 2880 resolution) using MATLAB (The MathWorks Inc.) and Psychtoolbox-3 (Brainard, 1997; Pelli, 1997). Face stimuli were generated using the Basel Face Model (Paysan et al., 2009), a morphable 3D face model derived from principal component analysis of high-resolution 3D face scans. Each face is defined by a vector of 199 shape coefficients, and linear combinations of these coefficients produce photorealistic face images spanning a continuous, high-dimensional face space. This representation allows the genetic algorithm to search a richly parameterized space of facial configurations without being constrained to a discrete set of pre-defined expressions.

### 2.4 Genetic Algorithm Face Task

Participants completed a genetic-algorithm face task in which they viewed a series of 7 charts, each containing 12 new faces, and selected any faces that manifest a target emotion. A genetic algorithm iteratively evolved faces to match their internal representation of each target emotion. The task was administered for 13 emotion categories: amusement, anger, awe, contempt, disgust, embarrassment, fear, happiness, interest, pride, sadness, shame, and surprise. The selection of these 13 categories was informed by Le Mau et al. (2021), who had professional actors portray emotional states in response to contextually rich scenarios; trained raters then evaluated these naturalistic expressions along 13 emotion dimensions, establishing an empirically grounded affective space that extends beyond traditional basic emotion taxonomies.

On each trial, participants viewed a population of 12 faces (each subtending approximately 6° of visual angle at a viewing distance of 60 cm, arranged in a 3 x 4 grid) and selected those that best matched the target emotion (e.g., ’Select the faces that look most embarrassed’). Participants could select between 0 and 12 faces per generation, with no time limit. Selected faces served as parents for the next generation: their 199-dimensional shape vectors were recombined via uniform crossover (each gene drawn from one of two randomly chosen parents with equal probability) to produce offspring. Half of the next generation’s faces (i.e. 6/12) were offspring of selected parents, and the other half were newly generated random faces. This process repeated for six selection generations (generations 1 through 6). In the seventh and final generation, the previously selected faces from the most recent generation(s) were re-presented, and participants chose the single face that best represented the target emotion. The order of emotions was randomized across participants.

### 2.5 Evolutionary Metrics

Five metrics were computed from each participant’s generational trajectory for each emotion, using data from all seven generations. For each generation, the mean of the 199-dimensional shape coefficient vector was computed, yielding a single scalar summary of the population’s position in face space at that generation. We refer to this value as the mean shape coefficient, denoted s. The five metrics were defined as follows.

**Intensity** was defined as the maximum s over the last three generations. This metric captures the peak strength of the evolved representation near the end of the search and serves as a measure of the *product* of face construction: what the participant ultimately produced, regardless of how they arrived there.

**Velocity** was defined as (s at generation 7 − s at generation 1) / number of generations. This metric captures the overall rate and direction of evolutionary change across the full trajectory.

**Stability** was defined as the standard deviation of s over the last five generations. Lower values indicate a more consistent search in which the population remained in a similar region of face space across generations; higher values indicate greater oscillation or drift.

**Range** was defined as the difference between the maximum and minimum s values across all seven generations. This metric captures the total breadth of the evolutionary search, reflecting how widely the participant explored face space.

**Convergence** was defined as the first generation at which s reached 90% of its maximum value across all generations. Earlier convergence (lower generation number) indicates faster crystallization of the representation. Unlike the four magnitude metrics, which are continuous measures derived from the shape coefficient values, convergence is an ordinal measure (generation number) that captures a qualitatively different construct: the *timing* of schema crystallization. This distinction motivated the separated multiple comparison correction described below.

### 2.6 Statistical Analysis

All analyses were conducted in Python 3.11 using NumPy, SciPy, and pandas, with all code available in a self-contained analysis script for reproducibility. Group comparisons were performed using Welch’s *t*-test, which does not assume equal variances and is robust to unequal sample sizes (Welch, 1947). Exact *p*-values were computed from the *t*-distribution using a two-tailed test.

Effect sizes were quantified as Cohen’s *d* computed with pooled standard deviations, with positive values indicating higher scores in the Low Risk group and negative values indicating higher scores in the At Risk group. Ninety-five percent confidence intervals for *d* were computed using the Hedges and Olkin (1985) variance approximation.

#### Multiple comparison correction

The 65 tests (13 emotions × 5 metrics) were divided into two families for Benjamini-Hochberg false discovery rate (FDR) correction: Family 1 comprised the 52 magnitude metric tests (4 metrics × 13 emotions), and Family 2 comprised the 13 convergence tests (1 metric × 13 emotions). This separation was motivated by the qualitative difference between the two measurement types: the magnitude metrics are continuous values derived from the shape coefficient space, whereas convergence is an ordinal generation number capturing a distinct construct (timing of schema crystallization). FDR correction was applied within each family at α = .05.

#### Permutation testing

To validate findings against violations of distributional assumptions, a label-shuffle permutation test was conducted for all results with uncorrected *p* < .10. On each of 5,000 iterations, group labels (Low Risk vs. At Risk) were randomly permuted and the Welch’s *t*-statistic recomputed, generating a null distribution against which the observed statistic was compared. The permutation *p*-value was defined as the proportion of null *t*-statistics equal to or more extreme than the observed *t*-statistic. All permutation tests used a fixed random seed for reproducibility (i.e., a deterministic starting state for the random number generator that ensures identical results on every rerun of the analysis).

#### Continuous analyses

To complement the group-based comparisons, Pearson and Spearman correlations were computed between continuous BDI-II total scores and each of the 65 evolutionary metrics. Spearman correlations were accompanied by 95% bootstrap confidence intervals (2,000 resampling iterations).

#### Sensitivity analyses

Three sets of sensitivity analyses assessed the robustness of results to analytic threshold choices. Convergence threshold sensitivity tested values of 80%, 85%, 90%, and 95% of peak. Stability window sensitivity tested windows of the last 3, 5, and 7 generations. Intensity window sensitivity tested windows of the last 2, 3, and 4 generations. For each parameter variation, the full analysis pipeline (Welch’s *t*-test, Cohen’s *d*, *p*-value) was recomputed.

#### Post-hoc power analysis

Achieved power for each observed effect size was estimated using the noncentral *t*-distribution with the observed Cohen’s *d*, sample sizes (*n* = 40 and *n* = 17), and α = .05 (two-tailed). The minimum detectable effect size at 80% power was also computed for the present sample sizes.

## 3. Results

### 3.1 Sample Characteristics

BDI-II scores ranged from 0 to 34 across the full sample (*M* = 10.42, *SD* = 7.44, Mdn = 9). The distribution was positively skewed, with 40 participants (70.2%) scoring in the minimal range and 17 (29.8%) meeting or exceeding the clinical cutoff of 14 (Figure 1). Within the At Risk group, 11 scored in the mild range (14–19), four in the moderate range (20–28), and two in the severe range (29–34). All 57 participants completed the GA face task for all 13 emotions with no missing data. Five quality checks were applied (missing BDI items, incomplete trial records, outlier detection, generation count verification, and data integrity), and no participants required exclusion on any criterion.

**Figure 1.**
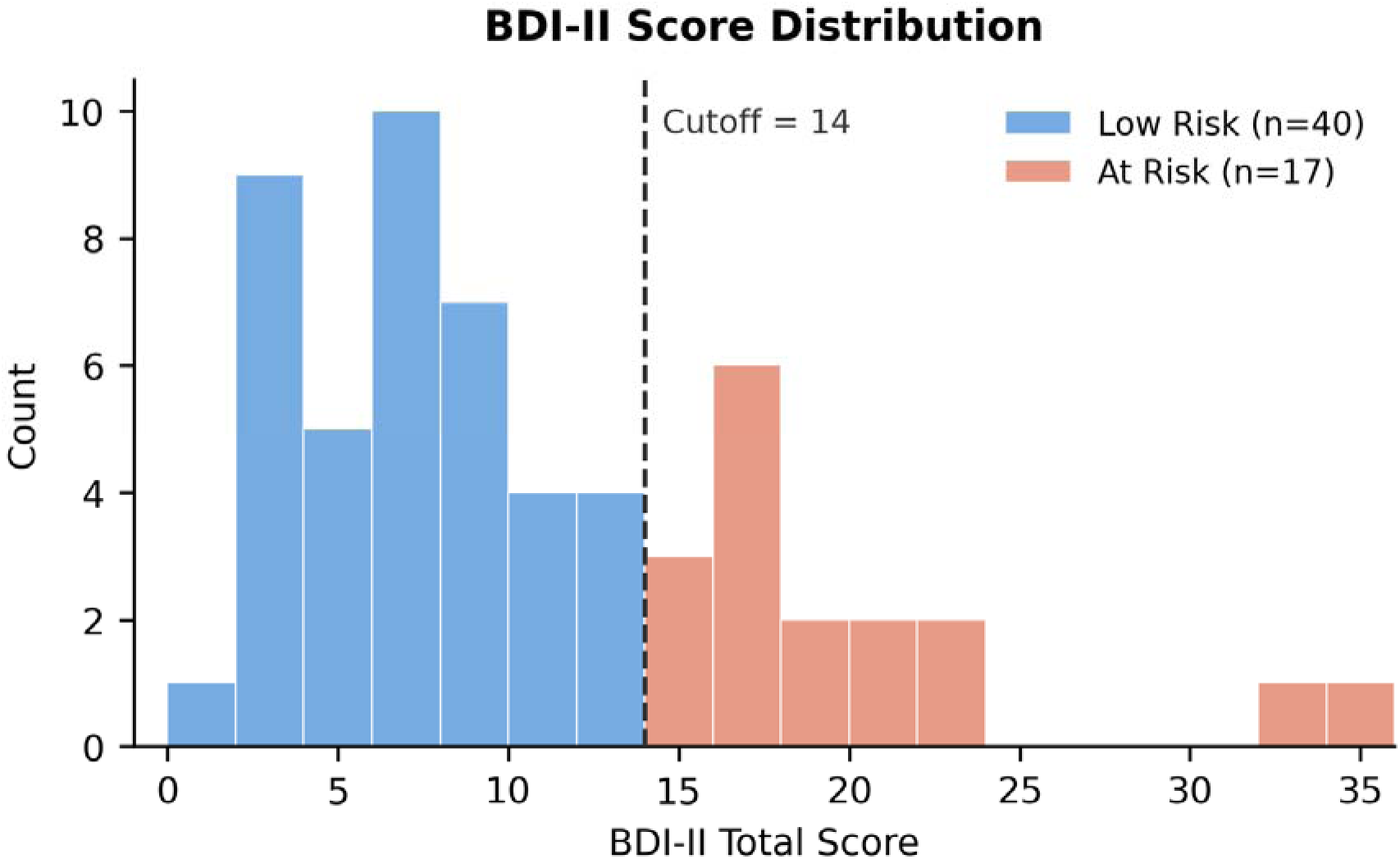
Distribution of Beck Depression Inventory-II (BDI-II) total scores across the sample. The dashed line indicates the clinical cutoff at 14, separating Low Risk (*n* = 40; blue) and At Risk (*n* = 17; coral) groups.

### 3.2 Primary Finding: Embarrassment Convergence

The strongest and most robust group difference was observed for embarrassment convergence. At Risk participants reached 90% of their peak embarrassment representation significantly earlier than Low Risk participants (At Risk: *M* = 1.71 generations, *SD* = 1.49; Low Risk: *M* = 3.45 generations, *SD* = 2.16), *t*(43.2) = 3.51, *p* = .001, Cohen’s *d* = 0.88, 95% CI [0.29, 1.47]. This result survived FDR correction within the convergence family (*p*FDR = .014) and was confirmed by permutation testing (*p*perm = .002). It was the only result in the entire analysis to survive FDR correction.

In practical terms, At Risk participants converged on their embarrassment template in roughly half the number of generations required by Low Risk participants. The evolutionary trajectories for embarrassment (Figure 2) illustrate this pattern: the At Risk group’s mean s plateaued or declined after an early peak, whereas the Low Risk group showed a more gradual, exploratory trajectory that continued to evolve across later generations.

**Figure 2.**
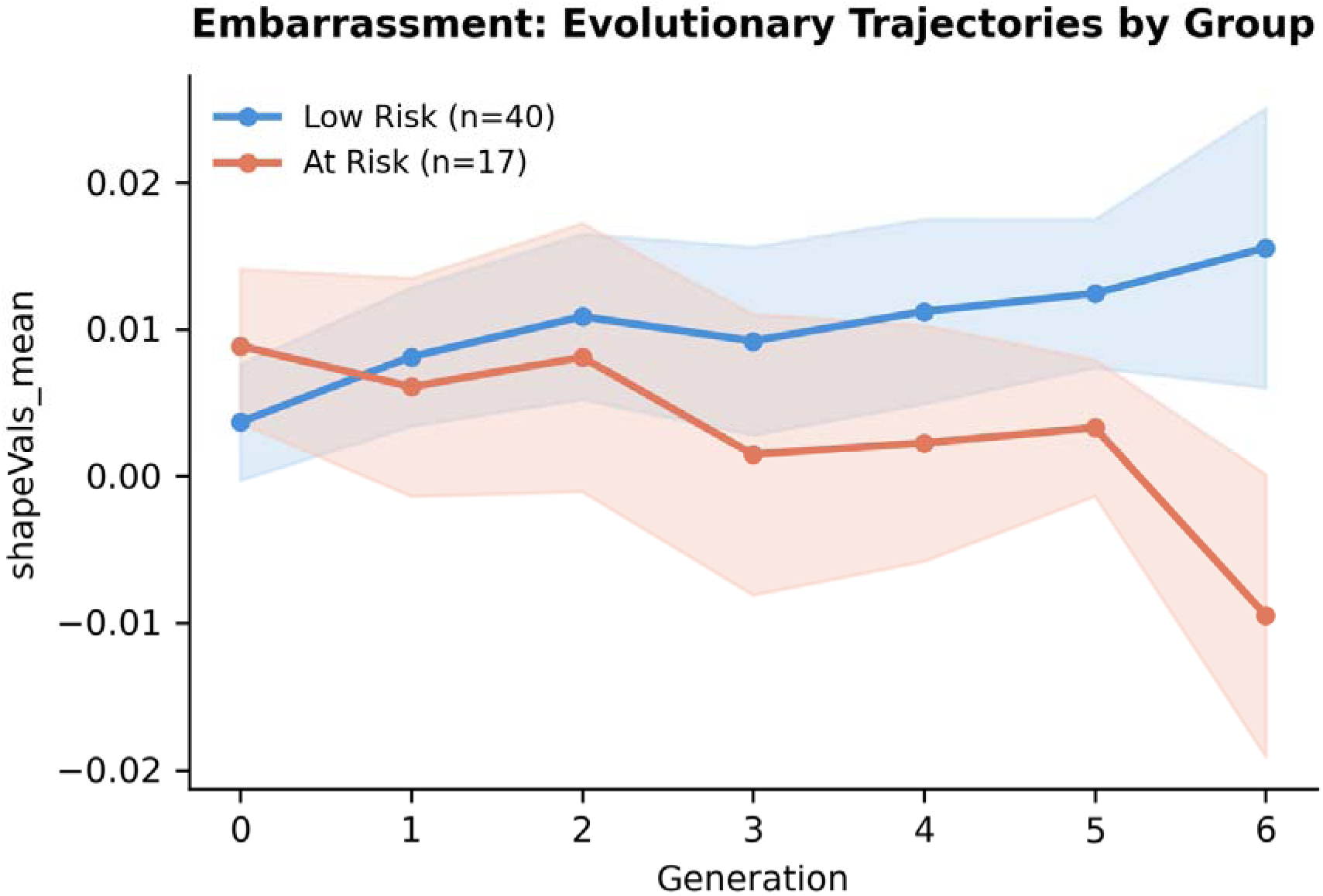
Evolutionary trajectories for embarrassment by group. Lines represent group means of the mean shape coefficient ( ) at each generation; shaded bands represent ±1 SEM. The At Risk group shows an early peak followed by decline, whereas the Low Risk group shows a more gradual, exploratory trajectory.

Sensitivity analyses confirmed that this finding was robust across all convergence threshold definitions. When the threshold was varied from 80% to 95% of peak, the effect size remained large (*d* = 0.80 to 0.93) and statistically significant at every threshold (all *p*s < .005; Figure 5). This robustness indicates that the group difference in embarrassment convergence is not an artifact of the specific threshold chosen.

### 3.3 Secondary Findings: Anger Metrics

Anger was the second emotion to show notable group differences, with effects emerging across multiple evolutionary metrics. However, none of the anger findings survived FDR correction, and they should therefore be interpreted with caution.

#### Anger stability

At Risk participants showed significantly greater variability in their anger representations across generations (At Risk: *M* = 0.029, *SD* = 0.013; Low Risk: *M* = 0.020, *SD* = 0.010), *t*(24.0) = -2.32, *p* = .029, *d* = -0.75, 95% CI [-1.34, -0.17], *p*perm = .030. The negative *d* indicates that the At Risk group’s anger trajectory was more unstable, oscillating more widely across generations rather than maintaining a consistent representation. Sensitivity analyses revealed that this effect was significant with a stability window of the last 5 generations (the primary analysis) but not with windows of the last 3 or 7 generations, indicating some dependence on the specific analytic window.

#### Anger intensity

At Risk participants also generated more intense anger faces (At Risk: *M* = 0.051, *SD* = 0.044; Low Risk: *M* = 0.025, *SD* = 0.041), *t*(28.4) = -2.08, *p* = .047, *d* = -0.62, 95% CI [-1.20, -0.04], *p*perm = .042.

#### Anger velocity

The rate of evolutionary change for anger showed a trending group difference in the same direction (*d* = -0.58, *p* = .068, *p*perm = .063), with At Risk participants showing a steeper upward trajectory across generations.

The anger trajectories (Figure 3) illustrate the pattern across these metrics: the At Risk group showed a strong, accelerating upward trend across generations, while the Low Risk group remained relatively flat near zero. This constellation of higher intensity, greater instability, and faster velocity in the At Risk group suggests a more volatile and less controlled anger search process.

**Figure 3.**
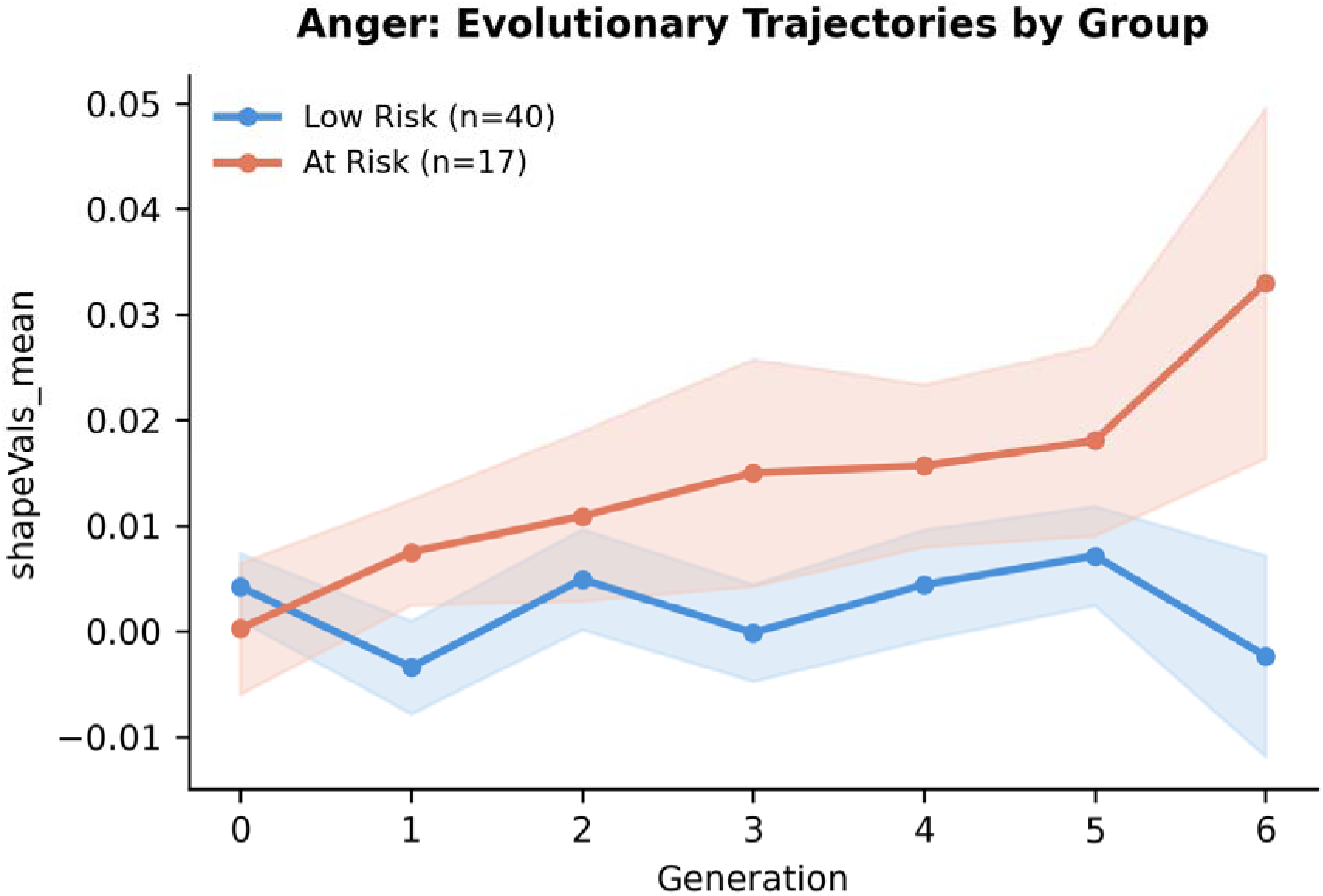
Evolutionary trajectories for anger by group. The At Risk group shows a strong, accelerating upward trend in across generations, while the Low Risk group remains relatively flat near zero. Shaded bands represent ±1 SEM.

### 3.4 Embarrassment Velocity

Embarrassment velocity also showed a significant group difference (*d* = 0.54, 95% CI [-0.04, 1.12], *p* = .033, *p*perm = .030), with At Risk participants showing a faster rate of evolutionary change for embarrassment faces. The positive *d* indicates that the Low Risk group showed higher (more positive) velocity values, suggesting a more sustained upward trajectory. This finding converges with the convergence result: At Risk participants not only arrived at their embarrassment template earlier but also showed a different pattern of evolutionary change across generations.

### 3.5 Overview of All Effects

Across all 65 tests, four reached uncorrected significance (*p* < .05) and were confirmed by permutation testing: embarrassment convergence, anger stability, embarrassment velocity, and anger intensity. An additional three tests showed trends at *p* < .10 (embarrassment intensity, anger velocity, contempt stability). The forest plot (Figure 4) displays the full distribution of effect sizes with 95% confidence intervals, and the effect size heatmap (Figure 6) provides an overview of all 13 emotions across all five metrics.

**Figure 4.**
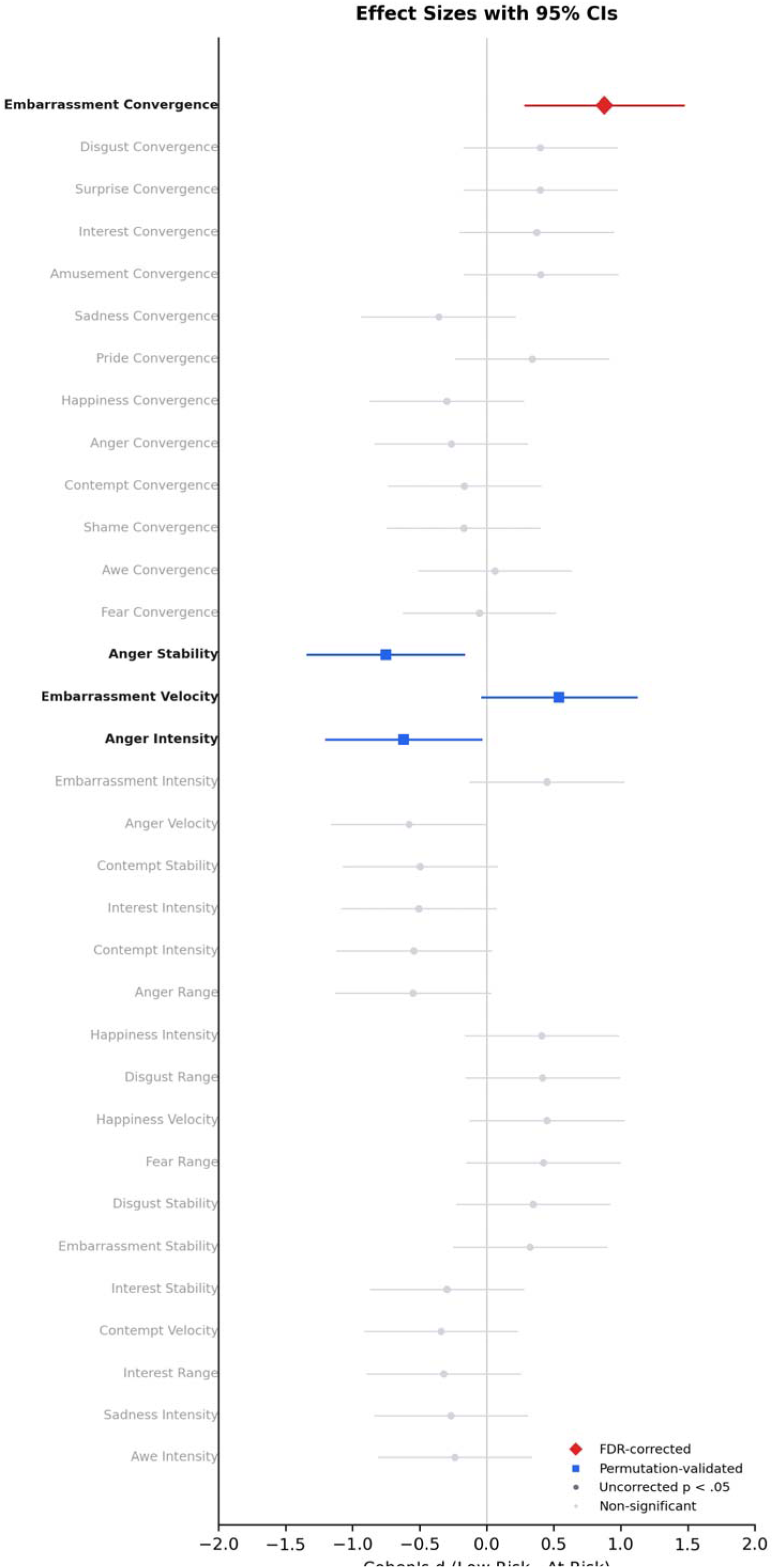
Forest plot of Cohen’s *d* with 95% confidence intervals for all group comparisons. Results are sorted by significance within two families: convergence metrics (top) and magnitude metrics (bottom). Red diamond = FDR-corrected significant; blue square = permutation-validated (*p*perm < .05); small purple diamond = uncorrected *p* < .05; gray circle = nonsignificant. Positive values indicate higher scores in the Low Risk group.

**Figure 5.**
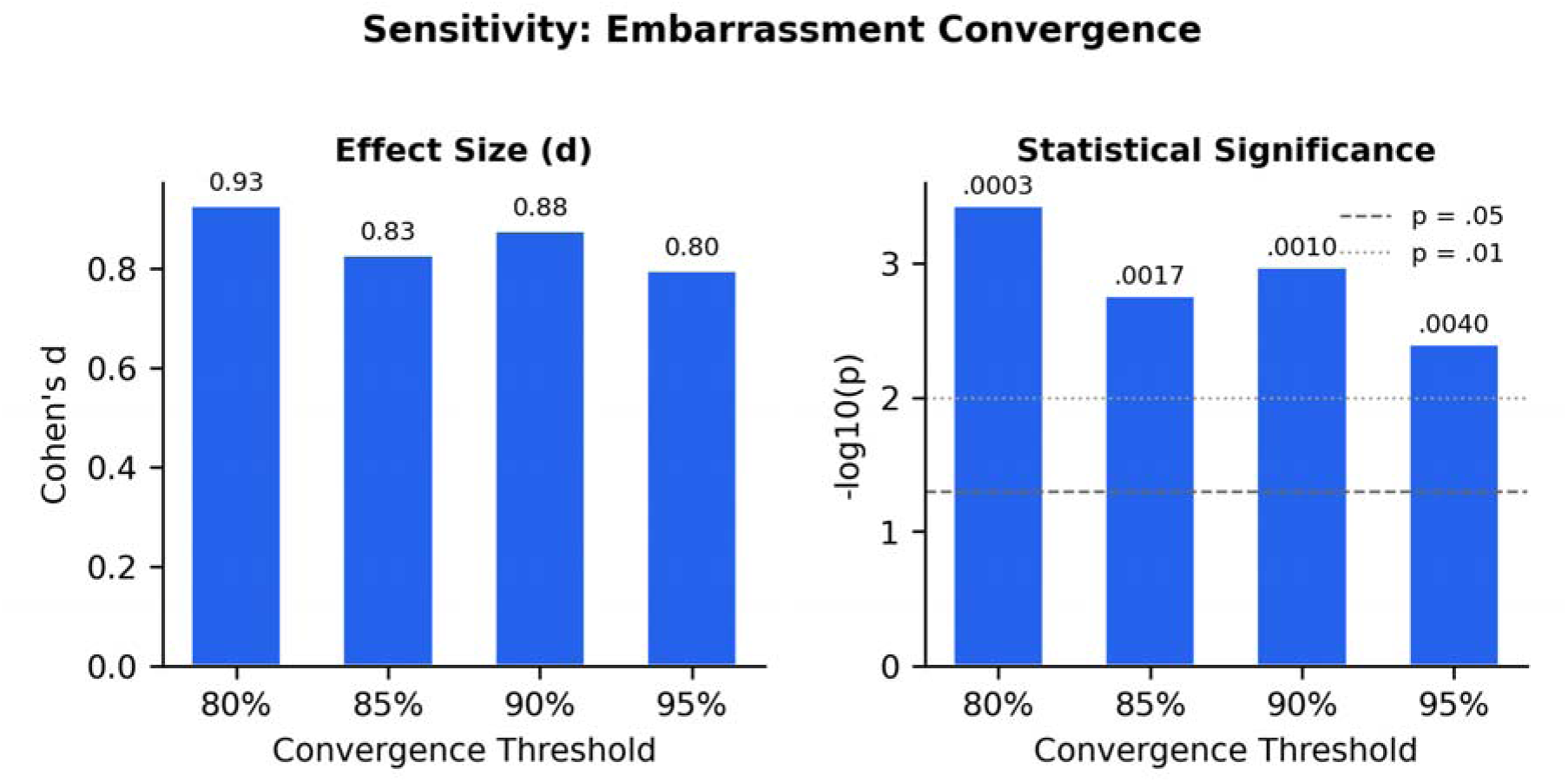
Sensitivity analysis for embarrassment convergence across four convergence threshold definitions (80%, 85%, 90%, 95% of peak). Left panel: Cohen’s *d* remains large (0.80 to 0.93) at all thresholds. Right panel: *p*-values (plotted as -log10) remain well below .01 at all thresholds. Dashed and dotted lines indicate *p* = .05 and *p* = .01, respectively.

**Figure 6.**
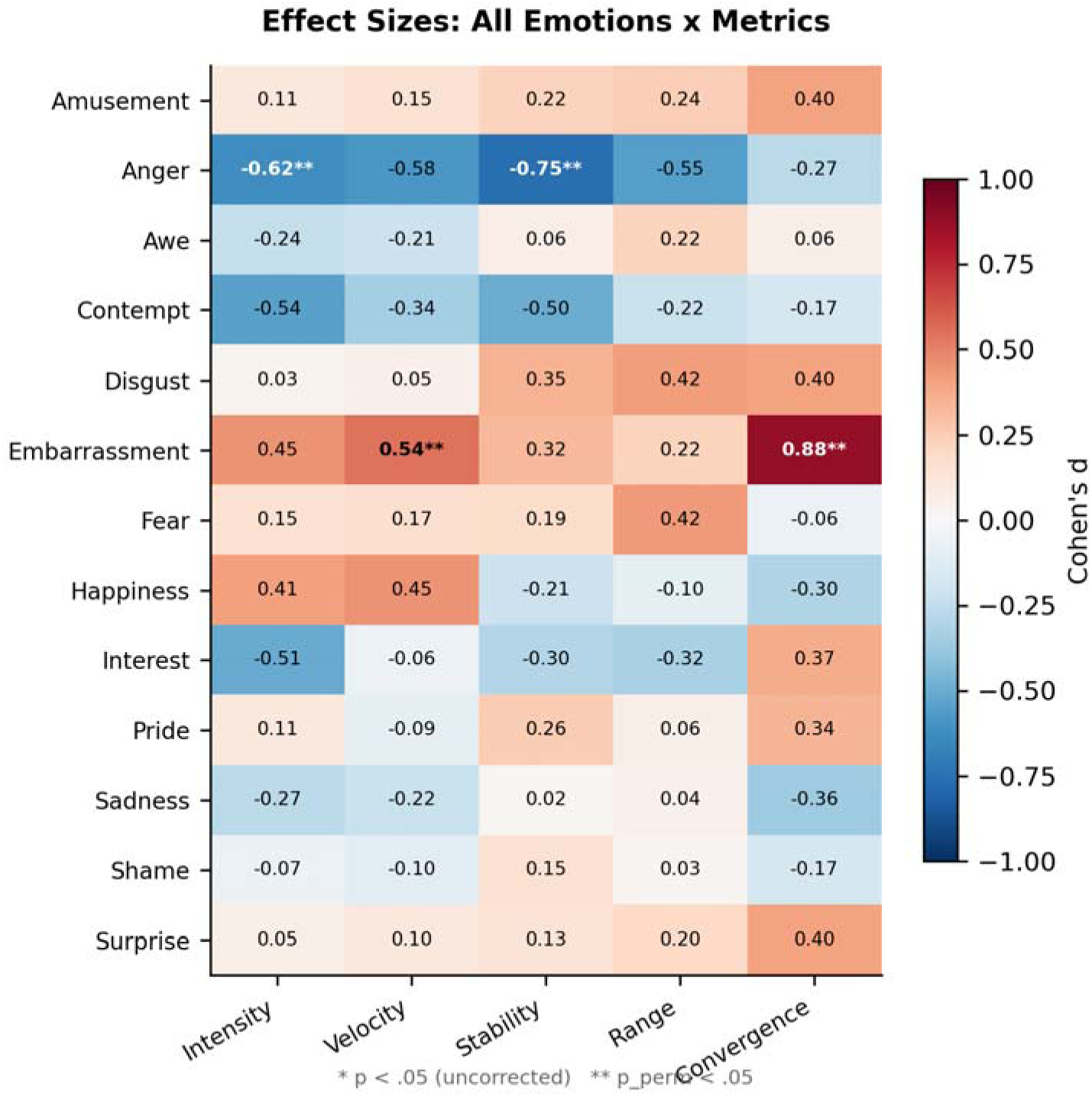
Effect size heatmap showing Cohen’s *d* for all 13 emotions across all five metrics. Warm colors (red) indicate higher values in the At Risk group; cool colors (blue) indicate higher values in the Low Risk group. Values are annotated with significance markers: * = uncorrected *p* < .05; ** = permutation-validated *p*perm < .05.

The overall pattern reveals that significant effects clustered around two emotions (embarrassment and anger) and were concentrated in process metrics (convergence, stability, velocity) rather than the product metric (intensity). The one exception, anger intensity, showed an effect in the opposite direction from the other intensity comparisons, which were uniformly nonsignificant.

### 3.6 The Product Null: Endpoint Intensities

A theoretically important pattern emerged in the intensity metric across emotions. With the exception of anger, none of the 13 emotions showed a significant group difference in peak intensity (all |*d*| < 0.55, all uncorrected *p*s > .06). The largest nonsignificant intensity effects were embarrassment (*d* = 0.45, *p* = .061) and contempt (*d* = -0.54, *p* = .102). Notably, sadness intensity showed essentially no group difference (*d* = -0.27, *p* = .354), nor did happiness (*d* = 0.41, *p* = .123) or fear (*d* = 0.15, *p* = .556). This pattern is consistent with the interpretation that depression modulates how participants construct their emotion representations (the process) rather than what they ultimately produce (the product).

### 3.7 Continuous BDI-II Correlations

Pearson and Spearman correlations between continuous BDI-II scores and the 65 evolutionary metrics yielded two uncorrected significant results. Fear range was negatively correlated with BDI-II (*r* = -.357, *p* = .006), and embarrassment convergence was negatively correlated with BDI-II (*r* = -.309, *p* = .019), indicating that higher depression severity was associated with earlier convergence on embarrassment representations. Neither correlation survived FDR correction. Pearson and Spearman results showed perfect concordance in significance across all 65 tests, indicating that the correlational findings were not driven by outliers or non-linear relationships.

### 3.8 Sensitivity Analyses and Power

Sensitivity analyses are summarized in Figure 5. The primary finding (embarrassment convergence) was robust across all tested parameter variations. When the convergence threshold was set at 80%, 85%, 90%, or 95% of peak, the effect size ranged from *d* = 0.80 to 0.93, and all *p*-values remained below .005. The anger stability finding was more sensitive to analytic choices: it reached significance with a stability window of the last 5 generations but not with the last 3 or 7 generations, though the direction of the effect was consistent across all windows.

Post-hoc power analysis indicated that the primary finding (*d* = 0.88) was adequately powered at 84.5%. The secondary anger findings, with effect sizes ranging from *d* = 0.58 to 0.75, had power estimates between 45% and 73%, indicating that the present sample was underpowered for detecting medium-sized effects. The minimum detectable effect size at 80% power for the current sample sizes (*n* = 40 and *n* = 17) was *d* = 0.83.

## 4. Discussion

### 4.1 Summary of Findings

The present study used a genetic-algorithm face synthesis task to examine whether depression severity modulates the process of constructing facial emotion representations. The central finding was a clear dissociation between process and product: At Risk participants differed from Low Risk participants in how they built their representations (the evolutionary dynamics of the search), but not in what they ultimately produced (the endpoint face intensity). This dissociation was emotion-specific, concentrated in embarrassment (a self-conscious emotion) and anger (an emotion linked to dysregulation in depression), rather than in the basic emotions most commonly studied in the depression and face perception literature.

The primary result was that At Risk participants converged on their embarrassment representations significantly faster than Low Risk participants, reaching their template in roughly half the number of generations. This was the only finding to survive FDR correction and was robust across all sensitivity analyses. A secondary cluster of effects emerged for anger, where At Risk participants showed greater evolutionary instability, higher peak intensity, and a trending increase in velocity. These anger effects were permutation-validated but did not survive FDR correction and were partially dependent on analytic window choices.

### 4.2 Embarrassment and Schema Rigidity

The accelerated convergence on embarrassment in the At Risk group is consistent with Beck’s cognitive schema theory (Beck, 2019). According to this framework, depression involves rigid, automatically activated negative schemas that bias information processing. In the context of the GA face task, schema rigidity would manifest as rapid deployment of a pre-formed template rather than open exploration of the face space. The At Risk group’s earlier convergence on embarrassment suggests that these participants possessed a more crystallized, readily accessible representation of embarrassment that was deployed quickly and with less generational exploration.

This interpretation aligns with the broader literature on self-conscious emotions and depression. Embarrassment requires the conjunction of self-evaluation and perceived social scrutiny (Keltner & Buswell, 1997), both of which are heightened in depression. The shame-depression link documented by Orth et al. (2006) suggests that self-conscious emotion schemas are particularly well-rehearsed in individuals with depressive symptomatology. Our finding extends this from the experiential domain (proneness to feeling embarrassed) to the representational domain (how an embarrassed face is constructed in a generative task). To our knowledge, this is the first evidence that depression shapes the generative process for self-conscious emotion faces specifically.

What does faster convergence mean in cognitive terms? In the GA face task, convergence speed reflects how quickly a participant’s selections become internally consistent—that is, how rapidly they identify and reliably select faces carrying the features they associate with the target emotion. Faster convergence thus implies that At Risk participants possessed a more crystallized embarrassment template that functioned as a stronger perceptual filter: they were quicker to detect embarrassment-relevant features in the randomly generated face populations and more decisive in selecting them, resulting in earlier stabilization of the evolutionary trajectory. This is consistent with a schema that is both more accessible (requiring less deliberation to activate) and more rigid (less open to alternative configurations of what embarrassment “looks like”). Importantly, faster convergence does not mean that At Risk participants produced different embarrassment faces; the endpoint intensity did not differ significantly between groups (*d* = 0.45, *p* = .061). The two groups arrived at similar representations of embarrassment, but the At Risk group got there faster and with less exploration—consistent with a pre-formed template that compressed the search process rather than a distorted endpoint.

### 4.3 Anger Instability and Emotion Dysregulation

The anger findings present a qualitatively different pattern from the embarrassment results. Rather than faster convergence (schema rigidity), the At Risk group showed greater instability in their anger trajectories: more oscillation across generations, higher peak intensity, and a steeper velocity of change. This pattern is consistent with the emotion dysregulation framework proposed by Besharat et al. (2013) and Loch et al. (2024), in which depression is associated with volatile and inflexible regulation of anger rather than a stable, well-consolidated anger schema.

The clustering of effects across three anger metrics (stability, intensity, velocity) strengthens the interpretive coherence of this finding, even though none of the individual effects survived FDR correction. The consistency of the direction across metrics suggests a genuine underlying pattern rather than isolated noise. Nevertheless, these results should be interpreted with appropriate caution. The anger stability effect was sensitive to the choice of stability window (significant at a window of 5 generations but not 3 or 7), indicating that the specific quantification of instability matters. Replication in a larger, adequately powered sample is needed before strong claims can be made about the anger findings.

Taken together, the embarrassment and anger results suggest that depression may have qualitatively different effects on different emotion categories. Self-conscious emotions, which are closely tied to the rigid self-evaluative schemas characteristic of depression, show a pattern of schema rigidity (faster convergence). Anger, which is linked to regulatory instability in depression, shows a pattern of dysregulation (greater trajectory volatility). This dual pattern is consistent with the view that depression involves both rigid negative cognitions and volatile emotional regulation (Gotlib & Joormann, 2010).

### 4.4 Why Not Sadness?

A notable null finding was the complete absence of group differences for sadness. Across all five metrics, sadness showed uniformly small effect sizes (all |*d*| < 0.27) and no significant differences. This extends the meta-analytic finding of Dalili et al. (2015), who reported that sadness recognition is the one emotion consistently preserved in MDD, from the domain of recognition to the domain of generation. Depressed individuals construct sadness faces no differently from non-depressed individuals, despite the centrality of sadness to the depressive experience.

One interpretation is that sadness representations are universally well-calibrated and not subject to schema distortion, perhaps because sadness is frequently encountered, socially reinforced, and culturally salient regardless of depressive status. An alternative interpretation is that the BDI-II cutoff, while valid for identifying elevated symptomatology, may not capture the severity threshold at which sadness processing becomes specifically altered.

### 4.5 The Mood Congruence Null

The continuous BDI-II correlations were weak and did not survive FDR correction. Only two of 65 correlations reached uncorrected significance, roughly what would be expected by chance at α = .05. This constrains mood-congruent processing theories: depression severity does not linearly amplify or distort emotion representations across the board. The effects observed in this study appear to be categorical (group-level schema differences) rather than dimensional (severity-scaled changes), suggesting that the relationship between depression and facial affect representation may have a threshold structure rather than a linear dose-response pattern.

### 4.6 Limitations

Several limitations should be acknowledged. First, the sample was subclinical: the At Risk group was defined by BDI-II scores at or above 14, a screening threshold that identifies elevated depressive symptomatology but does not constitute a clinical diagnosis of MDD. Although four participants in the At Risk group reported a prior depression diagnosis, no structured clinical interview (e.g., SCID, MINI) was administered. The term ’At Risk’ is used deliberately to avoid overinterpretation, and the findings should be understood as pertaining to dimensional variation in depressive symptoms rather than to clinical depression per se. This subclinical approach has limitations but also advantages: it avoids confounds associated with medication, hospitalization, and comorbidity, and is consistent with dimensional models of psychopathology (Gotlib & Joormann, 2010).

Second, the groups were unbalanced (*n* = 40 vs. *n* = 17). Although Welch’s *t*-test and permutation testing are robust to unequal sample sizes, the smaller At Risk group limits statistical power for detecting medium-sized effects. Post-hoc power analysis confirmed that the present sample was adequately powered only for large effects (*d* ≥ 0.83 at 80% power). The secondary anger findings, with effect sizes in the medium range (*d* = 0.58 to 0.75), were underpowered at 45–73%.

Third, the evolutionary metrics were computed from the mean shape coefficient (s), a scalar summary of the 199-dimensional face representation at each generation. This approach captures the overall magnitude of the evolved face in shape space but does not preserve information about the specific configuration of facial features. Future work could apply multivariate methods (e.g., cosine distance between generational populations, or representational similarity analysis across the full 199 dimensions) to examine whether depression modulates the direction of the evolutionary search in addition to its dynamics.

Fourth, the study was cross-sectional, precluding causal inferences. Schema rigidity for embarrassment could precede depressive symptoms (as a vulnerability factor), follow from them (as a consequence of repeated self-evaluative rumination), or both. Longitudinal designs are needed to distinguish these possibilities.

Fifth, the anger stability finding was sensitive to the choice of stability window, reaching significance with a window of the last 5 generations but not with windows of the last 3 or 7 generations. Although the direction of the effect was consistent across all windows, this analytic sensitivity warrants caution in interpreting the anger findings and underscores the need for replication.

### 4.7 Future Directions

Several directions emerge from these findings. First, replication in a clinical MDD sample with structured diagnostic interviews would test whether the embarrassment convergence effect amplifies with clinical severity or represents a subclinical-specific phenomenon. Second, longitudinal designs could examine whether schema rigidity for self-conscious emotions is a prospective risk factor for depression onset, which would have implications for early identification and prevention. Third, treatment studies could test whether cognitive-behavioral therapy (CBT), which explicitly targets rigid negative schemas, normalizes embarrassment convergence speed. If the effect reflects schema rigidity, it should be malleable through schema-focused interventions. Finally, a broader self-conscious emotion battery (adding guilt, regret, and social anxiety-related expressions) would clarify whether the embarrassment effect generalizes across the self-conscious emotion family or is specific to this category.

### 4.8 Conclusions

Depression severity modulates how people construct facial emotion representations, not what they ultimately produce. The genetic-algorithm face task revealed that individuals with elevated depressive symptoms converge faster on embarrassment representations (consistent with rigid self-conscious emotion schemas) and show greater instability in anger representations (consistent with emotion dysregulation). These effects were specific to self-conscious and anger-related emotions; basic emotions such as sadness, happiness, and fear showed no group differences on any metric. The findings extend cognitive schema theory to the domain of facial affect representation and highlight a previously unexplored dissociation between the process and product of emotion face construction. Self-conscious emotions, long recognized as central to depressive phenomenology, emerge here as a critical locus of depression-related perceptual bias.

